# *Crocus Sativus* and Its Active Compound, Crocin Inhibits the Endothelial Activation and Monocyte-Endothelial Cells Interaction in Stimulated Human Coronary Artery Endothelial Cells

**DOI:** 10.1101/2020.01.31.928341

**Authors:** Noor Alicezah Mohd Kasim, Nurul Ain Abu Bakar, Radzi Ahmad, Iman Nabilah Abd Rahim, Thuhairah Hasrah Abdul Rahman, Gabriele Ruth Anisah Froemming, Hapizah Mohd Nawawi

**Affiliations:** Department of Pathology, Universiti Teknologi MARA, 47000 Sungai Buloh, Selangor, Malaysia; Institute of Pathology, Laboratory and Forensic Medicine (I-PPerForM), Universiti Teknologi MARA, 47000 Sungai Buloh, Selangor, Malaysia; Faculty of Medicine, Universiti Teknologi MARA, 47000 Sungai Buloh, Selangor, Malaysia; Department of Basic Medical Sciences, Faculty of Medicine and Health Sciences, Universiti Malaysia Sarawak, Kota Samarahan, Malaysia

**Keywords:** *Crocus sativus*, Saffron, Crocin, Endothelial activation, Monocyte binding

## Abstract

*Crocus sativus L.* or saffron has been shown to have anti-atherogenic effects. However, its effects on key events in atherogenesis such as endothelial activation and monocyte-endothelial cell binding in lipolysaccharides (LPS)-stimulated *in vitro* model have not been extensively studied.

**Objectives:** To investigate the effects of saffron and its bioactive derivative crocin on the gene and protein expressions of biomarkers of endothelial activation in LPS stimulated human coronary artery endothelial cells (HCAECs).

**Methodology:** HCAECs were incubated with different concentrations of aqueous ethanolic extracts of saffron and crocin together with LPS. Protein and gene expressions of endothelial activation biomarkers were measured using ELISA and qRT-PCR, respectively. Adhesion of monocytes to HCAECs was detected by Rose Bengal staining. Methyl-thiazol-tetrazolium assay was carried out to assess cytotoxicity effects of saffron and crocin.

**Results:** Saffron and crocin up to 25.0 and 1.6 μg/ml respectively exhibited >85% cell viability. Saffron treatment reduced sICAM-1, sVCAM-1 and E-selectin proteins (concentrations: 3.13, 6.25, 12.5 and 25.0 μg/ml; 3.13, 12.5 and 25.0 μg/ml; 12.5 and 25.0, respectively) and gene expressions (concentration: 12.5 and 25.0μg/ml; 3.13, 6.25 and 25.0 μg/ml; 6.25, 12.5 25.0; respectively). Similarly, treatment with crocin reduced protein expressions of sICAM-1, sVCAM-1 and E-selectin (concentration: 0.2, 0.4, 0.8 and 1.6 μg/ml; 0.4, 0.8 and 1.6 μg/ml; 0.8 and 1.6 μg/ml; respectively] and gene expression (concentration: 0.8 and 1.6 μg/ml; 0.4, 0.8 and 1.6 μg/ml; and 1.6 μg/ml, respectively). Monocyte-endothelial cell interactions were reduced following saffron treatment at concentrations 6.3, 12.5 and 25.00 μg/ml. Similarly, crocin also suppressed cellular interactions at concentrations 0.04, 0.08, 1.60 μg/ml.

**Conclusion:** Saffron and crocin exhibits potent inhibitory action for endothelial activation and monocyte-endothelial cells interaction suggesting its potential anti-atherogenic properties.

## Introduction

Cardiovascular disease (CVD) is currently the leading cause of death in Malaysia with a percentage increase of 37.4% since 2007 (1). In Malaysia, CVD has been the leading cause among the five principals of morbidity and mortality; ischemic heart diseases (13.9%), pneumonia (12.7%), cerebrovascular diseases (7.1%), transport accidents (4.6%) and malignant neoplasm of trachea, bronchus and lung (2.3%) in 2017 (1,2). About 37 deaths per day due to CVD as compared to 24 deaths in 2007. Moreover, CVD is the principal cause of death among Malaysian men, aged 41 to 59 years old in urban areas (2). The major risk factors that cause CVD are smoking, high cholesterol level in blood, obesity and stress (3). The high death rate of atherosclerosis is owing primarily to the fact that it is a multifactorial disease. Therefore, research in recent years has very much skewed towards prevention of atherosclerosis.

Atherosclerosis is the underlying pathophysiology of CVD. Atherosclerosis was initially thought to be a degenerative disease which was an inevitable consequence of aging (4). Research in the last two decades, however, has shown that atherosclerosis is a multifactorial disease that encompasses both genetic and environmental factors (5). Atherosclerosis is a chronic inflammatory disease of the arteries, which is characterised by infiltration of leukocytes, deposition of lipids and thickening of vascular wall (6). Cellular and molecular events in the pathogenesis of atherosclerosis involves endothelial dysfunction due to an increase in endothelial activation biomarkers [soluble intercellular adhesion molecule-1 (sICAM-1), soluble vascular cellular adhesion molecule (sVCAM-1) and E-selectin] infiltration of monocytes, differentiation of monocytes into tissue resident macrophages and smooth muscle cell proliferation (7). The hallmark of atherosclerosis is the accumulation of cholesterol in the arterial wall, particularly cholesterol esters (8). Oxidised low-density lipoprotein (ox-LDL) plays an important role in the initiation and progression of atherosclerosis. The ox-LDL, recognised by scavenger receptors, is then taken up by the macrophages to form foam cells. These foam cells constitute a major source of secretory products that promote further progression of the atherosclerotic plaque (9).

Natural products that contain bioactive components, have been described to provide desirable health benefits beyond basic nutrition and are practically useful in the prevention of chronic diseases such as CVD and cancer (10). Saffron is a carotenoid-rich spice and also known as the ‘golden spice’ owing to its unique aroma, colour and flavour. It is derived from dried elongated stigma styles of a blue-purple flower, *Crocus sativus* L., traditionally used for several medicinal purposes such as a remedy for kidney problems, stomach ailments, depression, insomnia, measles, jaundice, cholera etc. in different parts of the world (11,12).

Four major compounds have been identified to be responsible in the health benefit profile of saffron. These compounds are crocin (colour), crocetin - central core of crocin, (colour), picrocrocin (taste) and safranal (aroma). Researchers have reported potential health benefits of these bioactive compounds (13). Crocin is a natural anti-oxidant with multi-unsaturated conjugate olefin acid structure. The compound exhibits favourable effects in the prevention and treatment of a variety of diseases which include dyslipidaemia, atherosclerosis, myocardial ischemia, haemorrhagic shock, cancer and arthritis (14). A study by Sheng et al., demonstrated inhibition of pancreatic lipase in rats by crocin (15). In addition, it has also been shown to inhibit formation of atherosclerosis in quails (16). These findings highlight the potential anti-atherogenic effects of crocin. However, studies to substantiate these effects at the cellular and molecular level remain scarce.

Therefore, this study aims to determine the role of crocin on atherogenesis at cellular level by examining its effects on the gene and protein expressions of biomarkers of endothelial activation and monocyte-cellular adhesion activity *in vitro.*

## Materials and Methods

### Cell culture

Human coronary artery endothelial cells (HCAECs) from Lonza, Switzerland were cultured in endothelial cell basal medium (EBM) supplemented with endothelial cell growth media (EGM) kits in 25cm^2^ flasks and incubated at 37°C in humidified 5% CO_2_ environment. The culture medium was replaced every 2 days until the cells were confluent. Cells with 80 to 85% confluency and only from passage 6 were used for the experiments.

### Preparation of crude extract

Saffron crude extract was prepared by dissolving 1g saffron dried filament (Sigma, Germany) into 10 ml of 75% ethanol and mixed thoroughly in supersonic water bath for 2 hours. The mixture then, was filtered and evaporated at 40°C and freeze dried to remove excessive ethanol and water (17). Saffron crude extract stocks of 0.2 g/ml and crocin (Sigma, Germany) stocks of 0.2 g/ml were dissolved in ethanol. The stocks were further diluted with treatment medium containing a mixture of 89% of RPMI-1640 (Sigma, Germany), 10% fetal bovine serum (FBS) (Gibco, USA) and 1% Streptomycin-penicillin to get working concentration of 200 μg/ml saffron crude extract and 200 μg/ml crocin with ethanol percentage less than 0.1%.

### Cell viability testing

Cell viability of HCAECs against saffron crude extract and crocin were tested using (3-(4, 5-Dimethylthiazol-2-yl)-2, 5-diphenyltetrazolium bromide (Invitrogen, USA), known as MTT assay. The HCAECs were seeded into 96 wells culture plate (1×10^4^ cells/well) and treated with various concentrations of saffron crude extract and crocin ranging from 1.6 to 400 μg/ml and 0.4 to 200 μg/ml respectively. The cells were incubated at 37°C for 24 hours in humidified 5% CO_2_ environment. Untreated cells were included as control wells. Then, 20 pl MTT solution (5 mg/ml MTT) was added to each well and incubated at 37°C for another 4 hours. The supernatant was then removed and replaced with 100 pl DMSO to dissolve the insoluble purple formazan product formed after previous incubation into a coloured solution, followed by 10 to 15 minutes incubation at room temperature. The absorbance was measured at 540 nm using a microplate reader (Tekan, Switzerland). The viability of the cells was measured by comparing the treated wells with the control wells and calculated using the following formula:

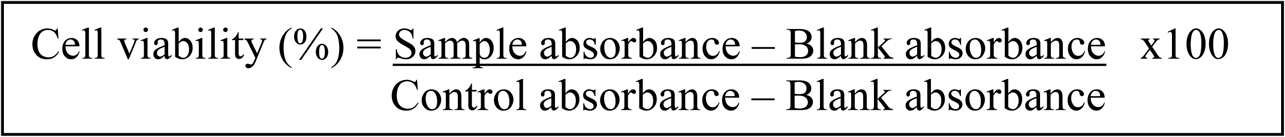

### Procarta cytokine analysis

Concentration of sICAM-1, sVCAM-1 and E-selectin were determined by measuring the protein expression for each respective biomarker in the supernatant of lipopolysaccharides (LPS)-stimulated treated HCAECs using Procarta Cytokine Assay Kit (Affimatryx, USA), bead based multiplex assay kit. All procedures were performed according to the manufacturer’s instruction and the signal is read using a Luminex instrument.

### Quantitative reverse transcription (qRT)-PCR analysis

HCAECs were harvested and extracted with RNeasy mini kit (Qiagen, USA). Concentration and purity of the total RNA was determined by Nanodrop and Agilent 2100 Bioanalyzer (Agilent, USA). Sensiscript reverse transcription kit (Bio-rad, USA) was used to reverse transcribe and amplify the RNA into cDNA. Oligonucleotide primers for ICAM-1, VCAM-1, E-selectin and glyceraldehyde 3-phosphate dehydrogenase (GAPDH) were purchased from Vivantis, USA while iQ™ SYBR® Green Supermix from Bio-rad, USA was used for quantitative assay. Real time PCR was performed on CFX96 in triplicates and normalised on the basis of their GAPDH content.

### Monocyte-endothelial interaction

HCAECs (Lonza, USA) were stimulated with LPS and treated with saffron and crocin extracts (Sigma, USA) at different concentrations ranging from 0.05 - 25.00 μg/ml and 0.001 - 1.600 μg/ml respectively. They were incubated for 16 hours. Monocyte U937 cell line (ATCC, USA) was added and incubated for 1 hour. 0.25% rose bengal was added and phosphate buffer solution (PBS) containing 10% FBS was used to remove the unbound cells. Ethanol:PBS (1:1) solution was used to stop the reaction. The absorbance was measured by spectrophotometer.

### Statistical analysis

All data were expressed as mean ± SD. Differences between groups were assessed by independent T-Test with SPSS (version 22, USA). The level of statistical significance was at *p* < 0.05.

## Results

### Effect of saffron and crocin on viability of HCAECs

**Fig 1(A)** demonstrates the viability of HCAECs following treatment with different concentrations of saffron ranging from 1.6 to 400.0 μg/ml, while **Fig 1(B)** shows the viability of HCAECs following treatment with different concentrations of crocin ranging from 0.4 to 200.0 μg/ml. Saffron and crocin concentrations of up to 25.0 μg/ml and 1.6 μg/ml, respectively gave more than 85% cell viability. Therefore, saffron concentration up to 25.0 μg/ml and crocin concentration up to 1.6 μg/ml was used for endothelial activation marker study.

**Fig 1.**
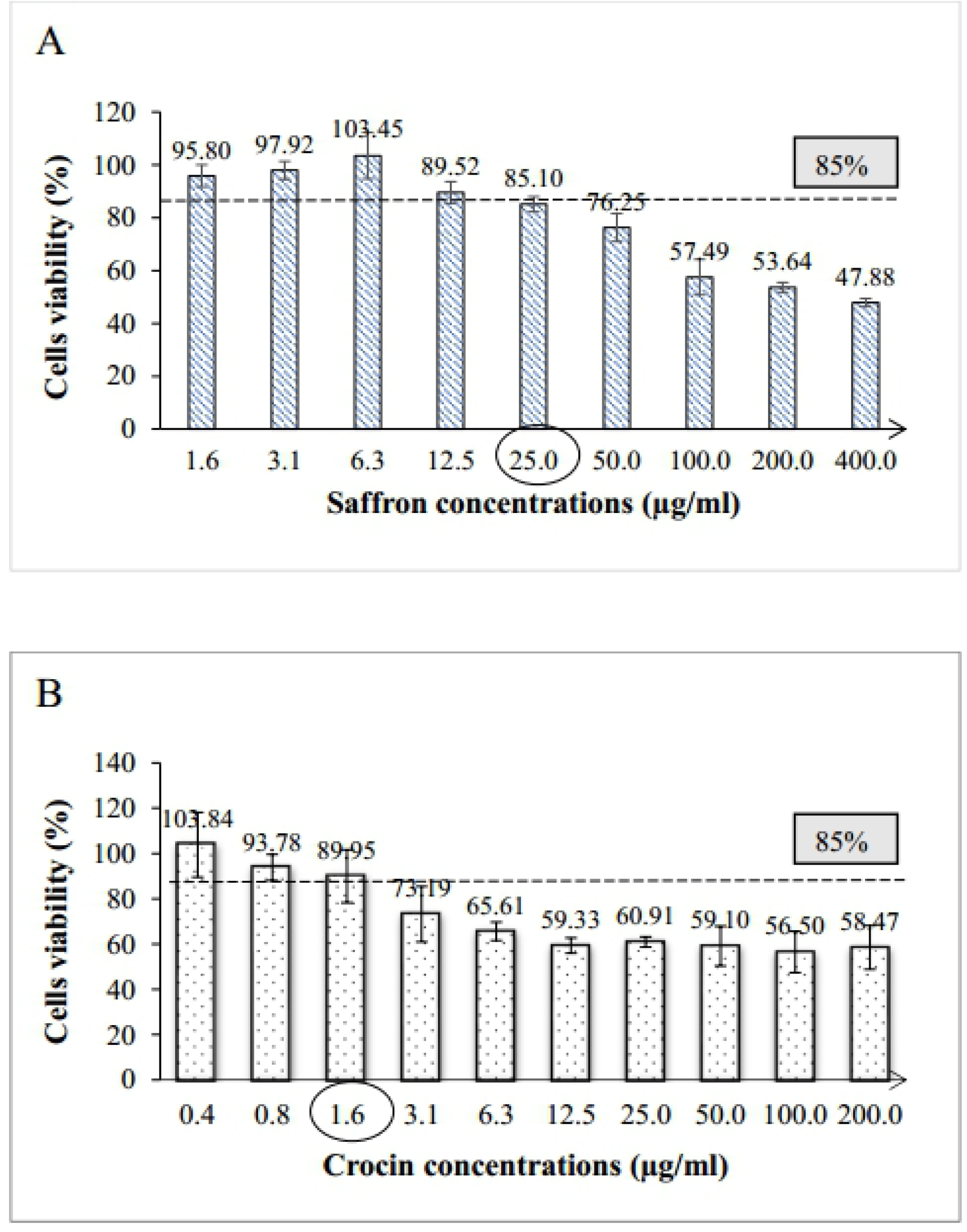
(A) Percentage (%) of human coronary artery endothelial cells (HCAECs) viability following treatment with various concentrations of saffron crude extracts (1.6¬400.0 μg/ml) and (B) crocin (0.4-200 μg/ml). Data are expressed as mean ± SD.

### Effects of saffron and crocin on protein expression of endothelial activation markers; sICAM-1, sVCAM-1 and E-selectin

ELISA kit was used to measure concentration of sICAM-1, sVCAM-1 and E-selectin expressed in the culture medium. The effects of saffron and crocin on endothelial activation markers were determined by comparing the concentration of each soluble molecule from treated LPS-stimulated sample with untreated LPS-stimulated sample. **Fig 2(A)** shows sICAM-1 levels significantly decreased following treatment of saffron and crocin at all doses [concentration: 3.13, 6.25, 12.5 and 25.0 μg/ml; 0.2, 0.4, 0.8 and 1.6 μg/ml. In **Fig 2(B)**, sVCAM-1 reduced following treatment with saffron at concentrations 3.13, 6.25 and 25.0 μg/ml (*p* <0.05) and crocin at 0.8 and 1.6 μg/ml (*p* <0.01). Untreated and non-stimulated sample served as negative control in this assay for validation. **Fig 2(C)** showed that E-selectin was reduced at higher doses of saffron (12.5 and 2.5 μg/ml) and crocin (0.4, 0.8 and 1.6 μg/ml) (*p* < 0.01).

**Fig 2.**
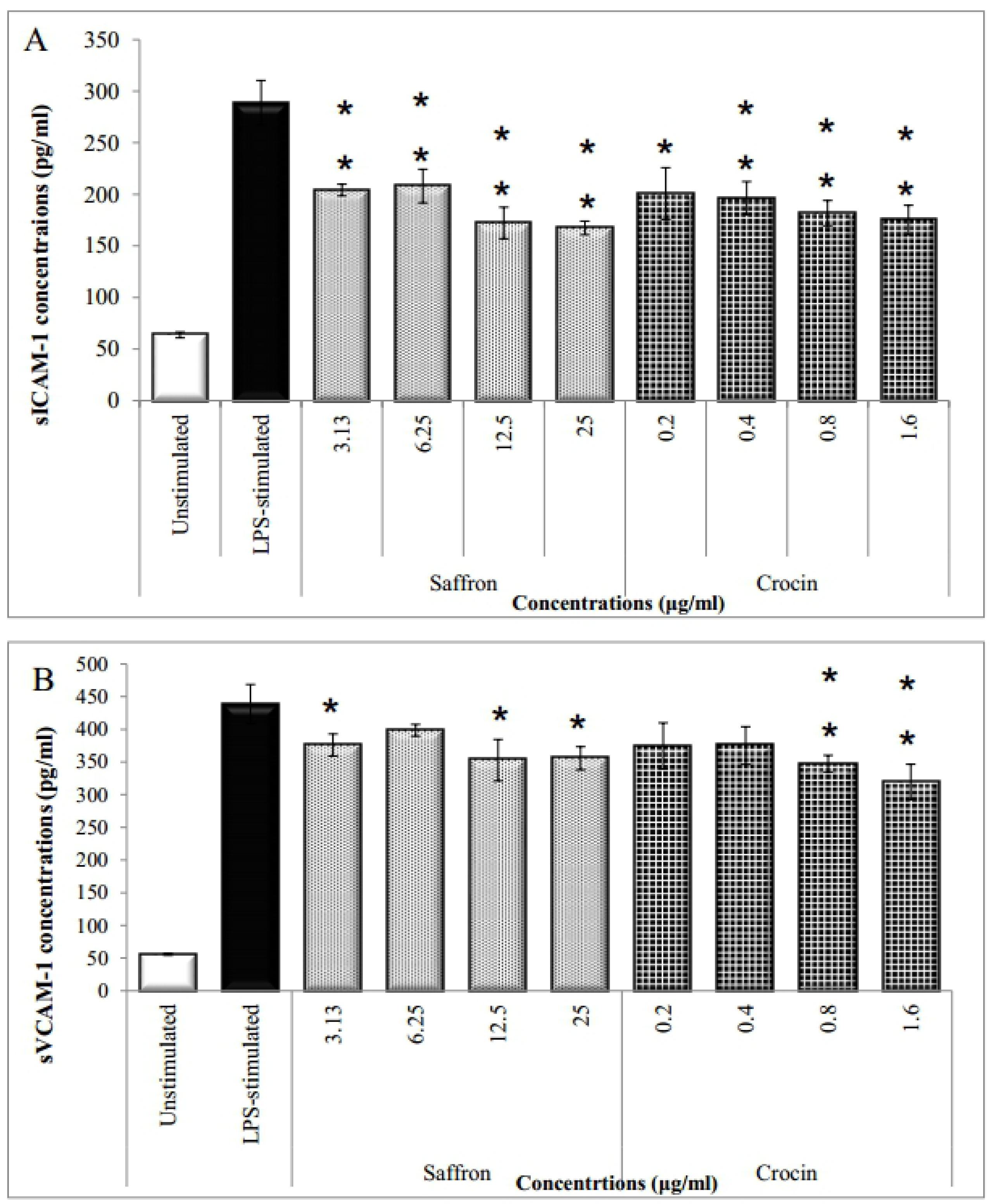
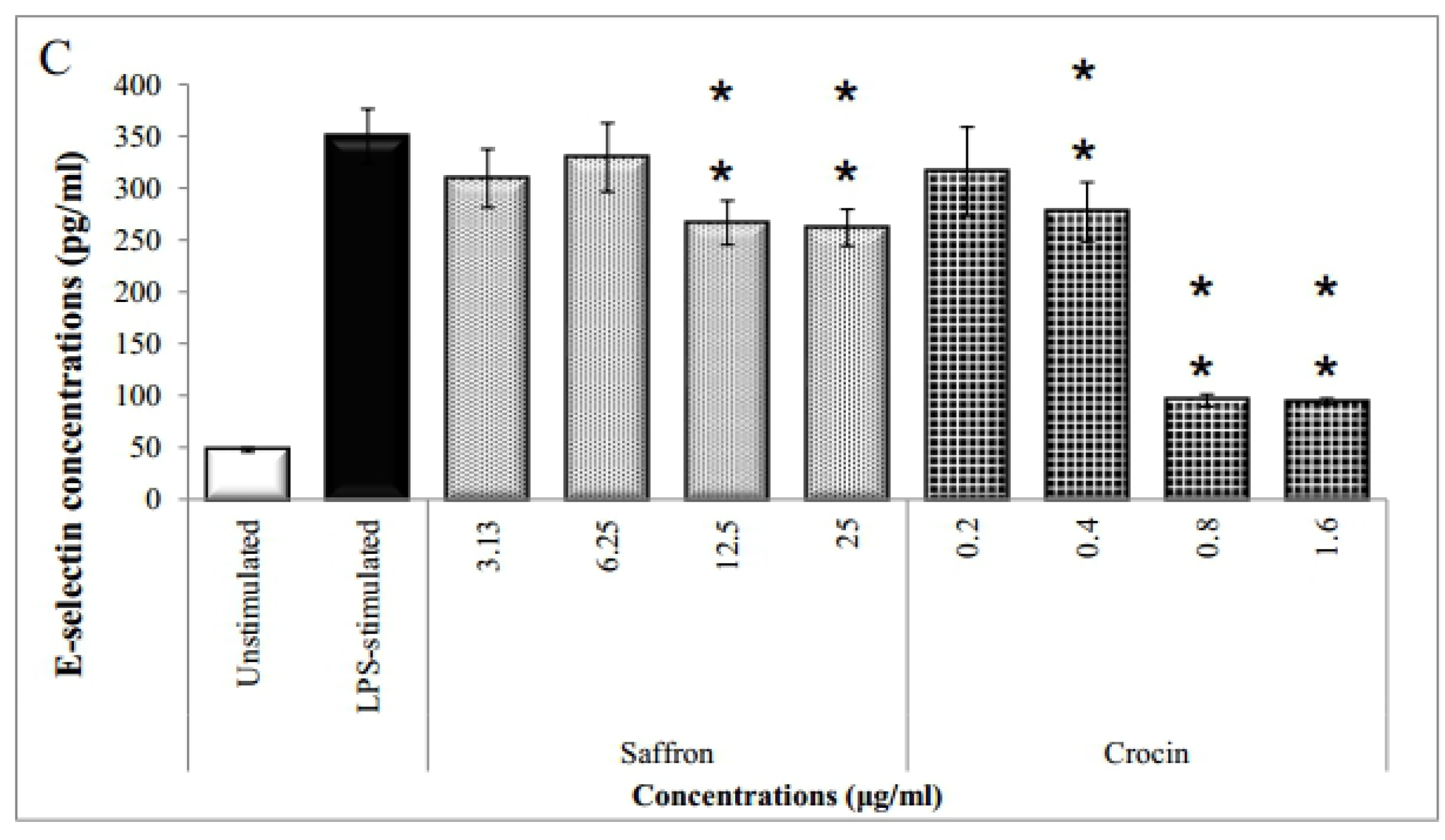
Effects of various concentrations of saffron crude extract (3.1.-25.0 μg/ml) and crocin (0.2-1.6 μg/ml) on (A) sICAM-1, (B) sVCAM-1 and (C) E-selectin protein expressions in HCAECs. Data are expressed as mean *±* SD. \**p* <0.05 and \*\**p* <0.01 compared to positive control. LPS; lipopolysaccharides, HCAECs; human coronary artery endothelial cells.

### Effects of saffron and crocin on gene regulation of endothelial activation markers; ICAM-1 VCAM-1 and E-selectin

CFX96 was used to measure ICAM-1, VCAM-1 and E-selectin gene regulation quantitatively from extracted HCAECS. Each sample was normalised to GAPDH gene and effects of saffron and crocin on gene regulation were determined by comparing treated samples with untreated LPS-stimulated sample. **Fig 3(A)** shows significant down-regulation of ICAM-1 gene after treated with saffron at doses of 12.5 and 25.0 μg/ml (*p* <0.05). Similarly, ICAM-1 gene reduced after treated with crocin at doses 0.8 and 1.6 μg/ml (*p* <0.05). In **Fig 3(B)** shows VCAM-1 genes were reduced at concentrations of 3.13, 6.25 and 25.0 μg/ml of saffron and 0.4, 0.8 and 1.6 μg/ml of crocin (\**p* <0.05 and \*\**p* <0.01). **Fig 3(C)** showed that E-selectin gene was significantly reduced at higher doses of saffron (6.25, 12.5 and 25 μg/ml) and crocin (1.6 μg/ml).

**Fig 3.**
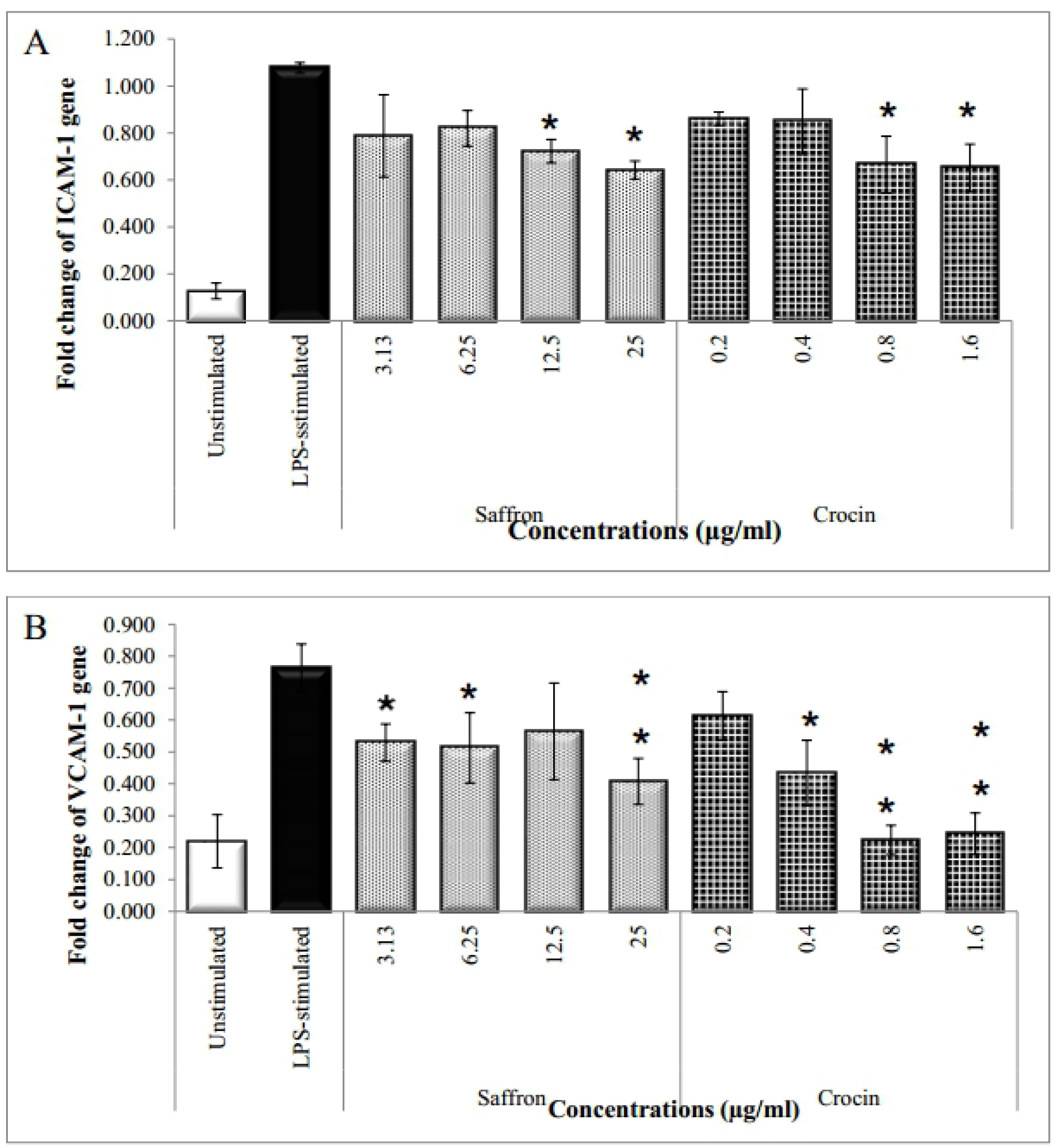
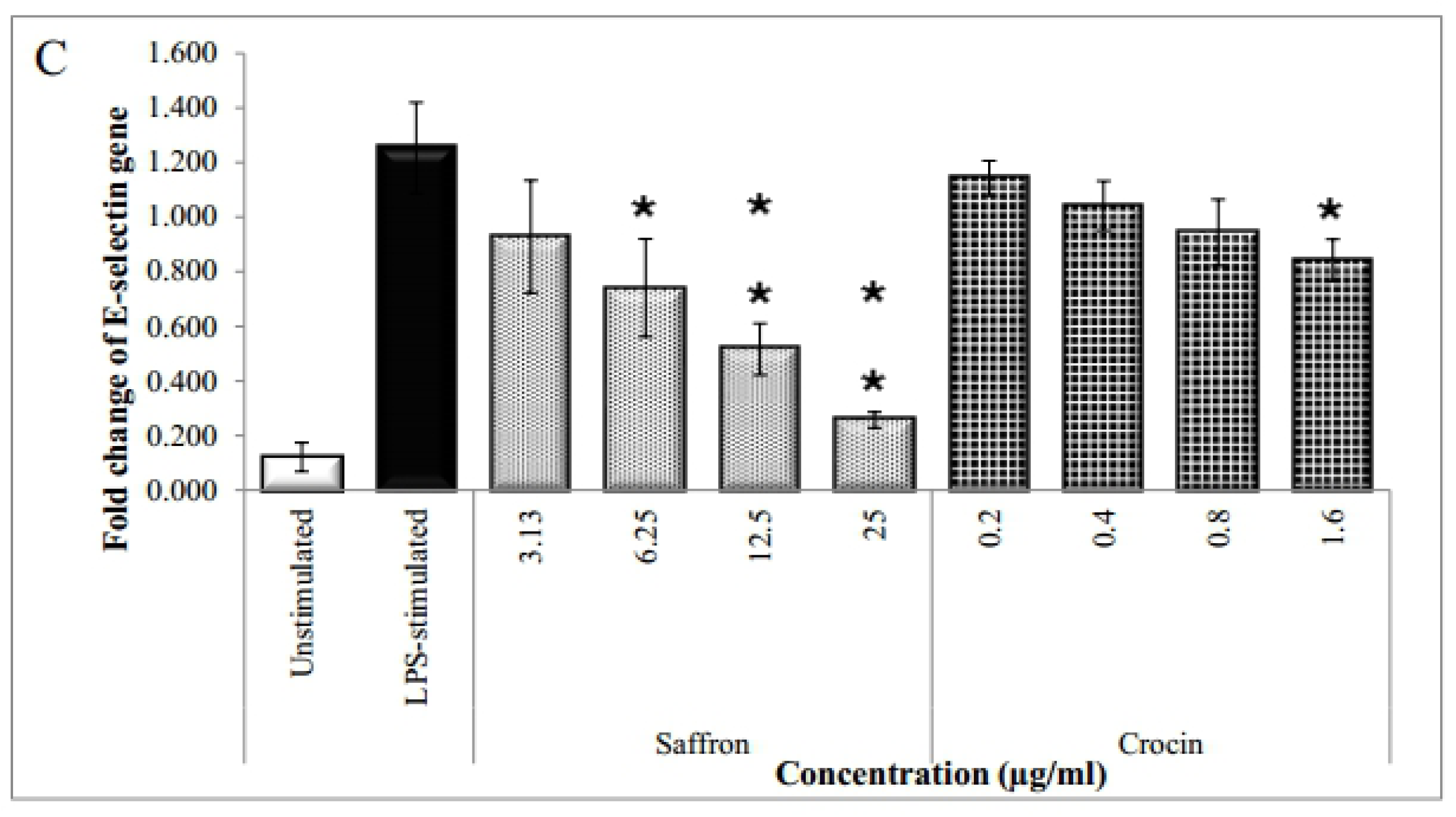
Effects of various concentrations of saffron crude extract (3.1-25.0 μg/ml) and crocin (0.2-1.6 μg/ml) on (A) ICAM-1, (B) VCAM-1 and (C) E-selectin gene regulation in HCAECs. Data are expressed as mean *±* SD. \**p* <0.05 and \*\**p* <0.01 compared to LPS-stimulated cells. LPS; lipopolysaccharides, HCAECs; human coronary artery endothelial cells.

### Effects of saffron and crocin on monocyte-endothelial cell interaction

Monocyte adhesion assay was performed to explore the effects of saffron and crocin on monocytes and endothelial cell interactions (**Fig 4**). Monocyte U937 cell line showed minimal adherence to the unstimulated HCAECs. After treatment with LPS for 16 hours, the adhesion of U937 monocytes to HCAECs was increased markedly. It was found that saffron and crocin at higher doses lead to the reduction of monocyte adhesion to LPS-stimulated cells. For saffron, significant reductions was observed at 6.25, 12.5 and 25 µg/ml (\**p* <0.05 and \*\**p* <0.01) compared to LPS-stimulated cells. Co-incubation of crocin at 0.4, 0.8 and 1.6 µg/ml significantly reduced the adhesion of monocytes to the LPS-stimulated HCAECs ml (\**p* <0.05 and \*\**p* <0.01).

**Fig 4.**
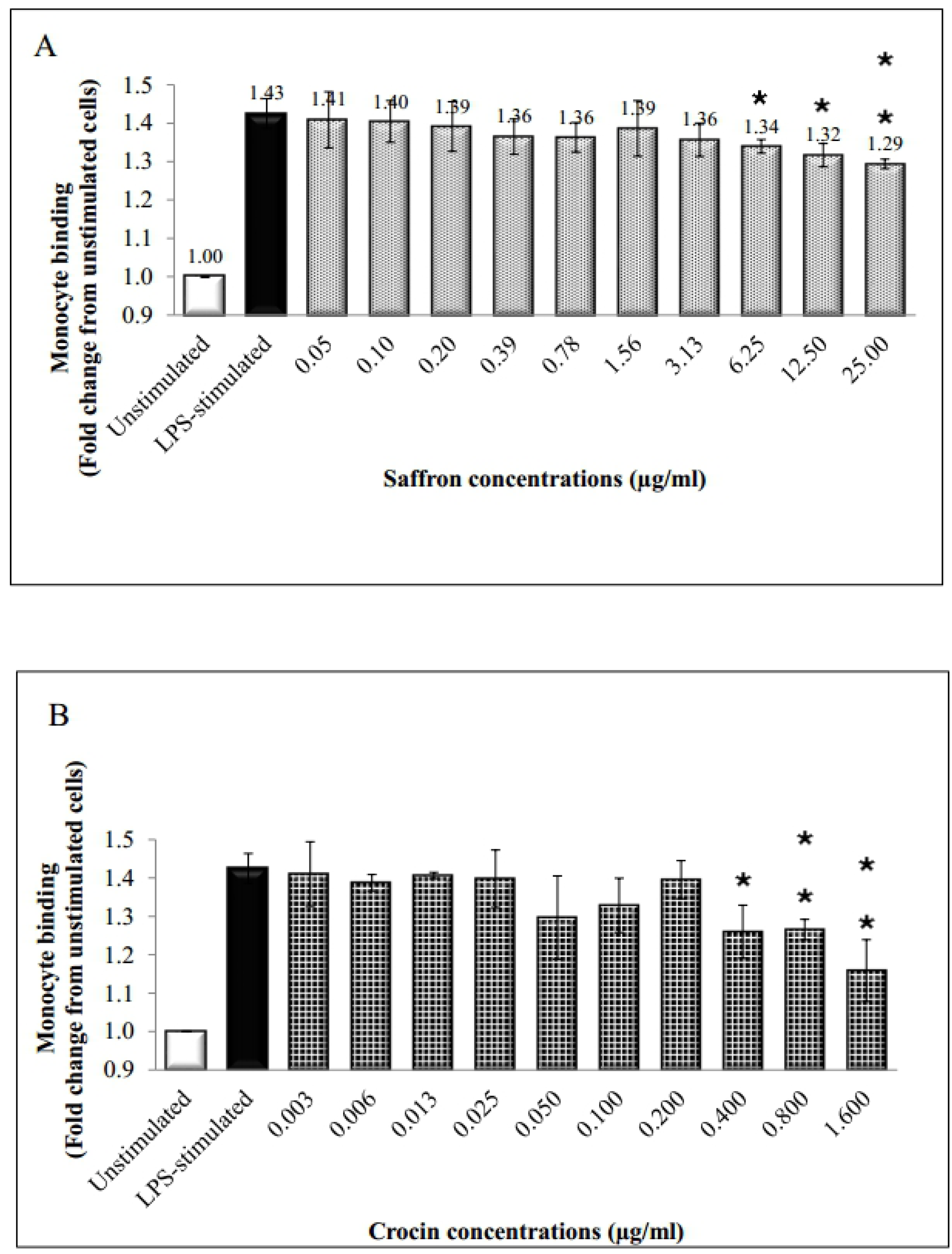
Effects of various concentrations of (A) saffron (0.05-25.00 μg/ml) and (B) crocin (0.003-1.600 μg/ml) on monocyte-HCAECs interaction. Data are expressed as mean *±* SD. \**p* <0.05 and \*\**p* <0.01 compared to LPS-stimulated cells. LPS; lipopolysaccharides, HCAECs; human coronary artery endothelial cells.

## Discussion

Studies on the effects of crocin on hyperlipidaemic animal models have demonstrated that it has potential anti-atherogenic properties by inhibiting pancreatic lipase and ox-LDL(15,16). At present, studies on anti-atherogenic effects of crocin mainly focused on ox-LDL related pathways. To the best of our knowledge, this is the first report to describe the anti-atherogenic effects of saffron and crocin at molecular and cellular levels. Although other works have published the inhibitory effects of crocin on ICAM-1 and VCAM-1 *in vitro*, its effect on endothelial markers in LPS-induced model has not been widely studied.

In the present study, LPS was used to activate an inflammatory response in endothelial cells to mimic the initial stage of atherosclerosis whereby inflammation precedes activation of endothelial biomarkers and accumulation of macrophages in tunica media region of artery(18). This study highlights the role of saffron crude extract and its bioactive compound, crocin, on the atherogenic pathway. Our group demonstrated a dose-dependent decrease in gene and protein expressions of biomarkers of endothelial activation (sICAM-1, sVCAM-1 and E-selectin) in LPS-stimulated HCAECs. This strongly suggests the role of saffron and crocin in attenuating atherogenesis. These results are consistent with the findings reported by other investigators using its active compound, crocin, which improved endothelial function via extracellular receptor kinase (ERK) and protein kinase B (or also known as Akt) signalling pathways in HUVECs(19).

sICAM-1 is an endothelial and leukocyte associated transmembrane protein renowned for its role in maintaining cell-cell interactions and promoting leukocyte endothelial transmigration. Activated endothelium releases soluble adhesion molecules and is therefore used to measure endothelial activation by measuring fluid-phase of those molecules(20). Previous study reported that another saffron compound, crocetin, can reduce leukocyte adhesion to endothelial cells by decreasing the expression of ICAM-1(18). This study shows that both saffron and crocin reduced the gene and protein expressions of sICAM-1, suggesting its potential role in attenuating monocyte transmigration.

A study by Zheng et al. (2005) reported that crocetin suppresses VCAM-1 expression due to the inhibition of nuclear factor-kappa beta (NF-κB) signaling, suggesting it’s anti-atherogenic effects in atherosclerotic-induced rabbits(21). Over expression of adhesion molecules, particularly VCAM-1, plays a central role in recruiting circulating monocytes into the intima as an initial event in the pathogenesis of atherosclerosis(22). Our study demonstrated a drop in gene and protein expressions of VCAM-1 which can be extrapolated to denote that recruitment of monocytes into the intima may be inhibited.

Likewise, E-selectin is a carbohydrate-binding molecule located in endothelial cells, in which it is responsible for the attachment and gradual rolling of leucocytes in an inflammatory response along the vascular wall(23). There is also evidence that the interaction of selectins (E-selectin, P-selectin, L-selectin) with glycosylated ligands of leucocytes mediate rolling and also their behaviour by enabling integrin-dependent reductions in rolling velocities. Selectins also mediate leucocyte adhesion to activated platelets and to other leucocytes. Such results showed that selectins initiate multicellular adhesive and signalling events during pathological inflammation(24). Thus, selectins are known to be endothelial activation biomarkers.

Recruitment of monocytes is also an important event in the process of atherosclerotic plaque formation. This process is influenced by chemo attractants, adhesion molecules and some receptors. Monocytes differentiate into macrophages in the vessel wall and start to take up lipids, which result in the transformation of macrophage into foam cells^30^. Scarce data have been reported on the effects of saffron and crocin on monocyte adhesion. In the present study, the unstimulated group showed minimal binding to monocyte U937 cells, whereas co-incubation with LPS markedly increased the adhesion of monocytes to HCAECs. This study revealed that saffron and crocin could inhibit monocytes adhesion to endothelial cells at higher concentrations. The increased potency of saffron and crocin is likely to be due to greater beneficial effects in terms of inhibition of adhesion molecules.

This study demonstrates that saffron crude extract and its bioactive compound, crocin are potent anti-atherosclerotic agents for the suppression of endothelial activation biomarkers by reducing the mRNA levels of ICAM-1, VCAM-1 and E-selectin thereby inhibiting its protein synthesis. This in turn explains the reduction in monocyte adhesion as these biomarkers are significant in the recruitment and transmigration of monocytes to the tunica intima. However, saffron crude extracts exhibit better anti-atherosclerotic properties by reducing E-selectin effectively compared to crocin. This may be due to the synergistic effects of other bioactive compound found in the crude extract(25). These findings suggest that, although both compounds show substantial anti-atherogenic properties, saffron appears to exhibit more prominent effect, possibly owing to other bioactive compounds in the crude extract that collectively exerts a more significant anti-atherogenic effects. Future studies to identify other bioactive compounds from crude saffron and determine their effects on atherogenesis would further ascertain the potential of saffron as an anti-atherogenic supplement. In addition, *in vivo* studies on crocin and its bioactive compound would determine if in fact, these compounds can be a potential addition to current treatment modalities to the prevention of CAD.

## Conclusion

Saffron and crocin exhibits potent inhibitory action for endothelial activation, suggesting its potential anti-atherogenic properties where saffron shows better reducing effects on E-selectin. Both compounds also reduce monocyte-endothelial cell interaction. However, saffron exerts more inhibitory effect, suggesting its effectiveness as anti-atherogenesis than its active compound, crocin.

## Acknowledgement

The authors thank Institute for Medical Molecular Biotechnology (IMMB), Faculty of Medicine, Universiti Teknologi MARA for providing necessary facilities.

